# Diurnal Regulation of Urinary Behavior and Gene Expression in Aged Mice

**DOI:** 10.1101/2025.03.16.642675

**Authors:** Danielle S. Taylor, Albert A. Allotey, Rachel E. Fanelli, Sushumna B. Satyanarayana, Sharanya S. Bettadapura, Cole R. Wyatt, Jason G. Landen, Adam C. Nelson, Emily E. Schmitt, Danielle R. Bruns, Nicole L. Bedford

## Abstract

Nocturia, defined as waking one or more times per night to urinate, is a prevalent and burdensome condition with few effective treatments. While the primary risk factor for nocturia is advanced age, few preclinical studies have addressed the pathophysiological mechanisms of nocturia in older subjects. Here, we develop a translational model of nocturia using aging mice and a behavioral paradigm that enables circadian assessment of voluntary urination in group-housed animals. We discovered dampened diurnal regulation of urinary behavior in aged mice compared to adult controls. Molecular analyses revealed disrupted diurnal expression of canonical circadian genes in aged mouse kidney and bladder tissues. Notably, we identified age-related loss of diurnal regulation of the bladder mechanosensory ion channel, *Piezo*1, suggesting a potential mechanism linking circadian disruption to altered bladder sensitivity. Our results reveal a role for circadian dysfunction in age-related nocturia and identify *Piezo*1 as a promising therapeutic target for chronobiological intervention.

## INTRODUCTION

Across mammals, including humans, urination predominantly occurs during the active phase—a trend that persists even under constant darkness (1, 2). Nocturia, the need to wake during the night to urinate, is one of the most common and troublesome urinary conditions. While it affects people of all ages, nocturia primarily impacts older adults, with approximately 70% of individuals over age 60 reporting symptoms (3, 4). In this population, nocturia is associated with increased risk of falls and fractures, reduced sleep quality, and numerous comorbidities including cardiovascular disease (5-7). Current therapeutics are poorly tolerated and largely unsuccessful, due in part to limited understanding of the underlying pathophysiology and molecular mechanisms of age-related nocturia (8, 9). Additionally, although nocturia specifically refers to dysregulated voiding at night (10), it is rarely studied as a circadian disorder.

Circadian rhythms play a critical role in human health (11). Consequently, their disruption may result in maladaptive health outcomes, including dysregulated urination. For example, disrupted clock gene expression through aging or shift-work is associated with renal and urinary system dysfunction (12, 13). Moreover, global deletion of *Clock* results in a nocturia-like phenotype in young male mice (14), suggesting a fundamental link between the circadian system and urination patterns. We therefore hypothesize that circadian disruption in healthy aging contributes to nocturia. However, whether laboratory mice—a widely used animal model of human aging (15, 16)—exhibit age-related circadian dysregulation of urination behavior remains unclear.

To address this gap, we developed a behavioral paradigm that quantifies daily urinary behavior in group-housed mice. Capitalizing on the natural tendency of mice to segregate their nesting and elimination sites (17), we designed a 48-hour video recorded latrine cage assay that avoids the chronic stress associated with traditional metabolic cages (18, 19). Using this approach, we examined circadian rhythms of locomotor and urination behavior in aged mice and adult controls. We also analyzed age-specific expression of canonical circadian genes in key urinary system organs. In addition, we investigated the effect of aging on diurnal expression of the bladder mechanosensory gene, *Piezo*1, which may contribute to daily variation in functional bladder capacity (20). Together, understanding how age-related circadian dysregulation contributes to nocturia at a behavioral and molecular level will offer a novel scientific framework for developing chronotimed therapies.

## METHODS

See Supplementary Material for full methodological details.

## RESULTS

### Circadian Rhythm of Core Body Temperature (Tb) is Dampened in Aged Mice

To assess age-related changes in circadian rhythms, we measured daily core body temperature (Tb) in adult (4-5 months) and aged (19-21 months) mice of both sexes (**Fig. 1A**). To determine how precisely the Tb rhythm aligned with the 24-hour light-dark cycle, we generated chi-square periodograms in ClockLab (Actimetric) and used linear models (LMs) to test for differences in period and amplitude (**Fig. 1B**). While there was no effect of age or sex on period (LM, age: *t* = 0.393, *P* = 0.700; sex: *t* = 0.321, *P* = 0.753, age*sex: *t* = 0.321, *P* = 0.752), we observed a significant effect of both age and sex on amplitude (**Fig. 1C**; LM, age: *t* = 2.890, *P* = 0.011; sex: *t*= 3.240, *P* = 0.005, age*sex: *t* = 0.577, *P* = 0.573). This effect was primarily driven by females, with aged mice showing significantly dampened circadian Tb rhythms compared to adults (**Fig. 1C**; LMs, female: *t* = 5.670, *P* < 0.001; male: *t* = 1.664, *P* = 0.135).

**Figure 1.**
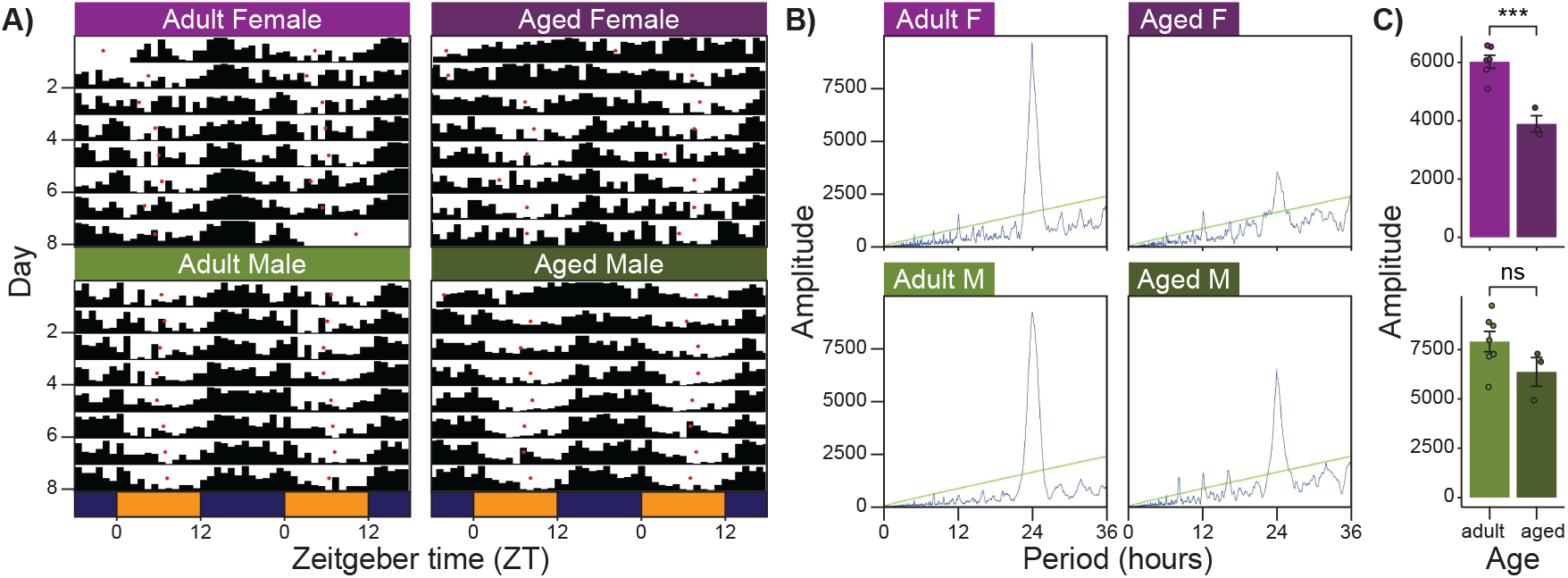
Circadian rhythms of core body temperature (Tb) in adult and aged mice. Representative Tb actograms (**A)** and periodograms **(B). C)** Differences in Tb amplitude between adult and aged females (top) and males (bottom). Error bars represent SEM. Significance: \*\*\**P* <0.001. Sample size: *n* = 6 adult female, 3 aged female, 7 adult male, and 3 aged male mice.

### Diurnal Regulation of Locomotor Activity is Dampened in Aged Mice

When given access to multiple cages, mice will segregate their living and toileting areas, preferentially urinating in one cage (17). We leveraged this natural behavior to non-invasively quantify daily urination patterns using a latrine cage assay (LCA). Specifically, we adapted a complex housing system where two cages connected by an external tunnel serve distinct purposes: a *home cage* (containing food, water, bedding, nestlet, and red hut) and a *latrine cage* lined with custom-cut filter paper (**Fig. 2A**). Filter papers were exchanged every 12 hours and imaged under blue light to quantify urine cover. We quantified locomotor activity as frame-to-frame pixel changes and compared activity between the light and dark phases in both cages (**Fig. 2B**). Both age groups spent the majority of their time in the home cage (73% on average). Using linear mixed-effects models (LMMs) that accounted for the number of mice per cage (fixed effect) and cage ID (random effect), we found significantly higher total cumulative activity (home + latrine cage) during the dark phase —which, in nocturnal mice, represents the active phase—across all four groups (**Fig. 2C**; LMMs, adult female: *t* = 8.603, *P* < 0.001; aged female: *t* = 8.550, *P* < 0.001; adult male: *t* = 16.591, *P* < 0.001; aged male: *t* = 10.718, *P* < 0.001).

**Figure 2.**
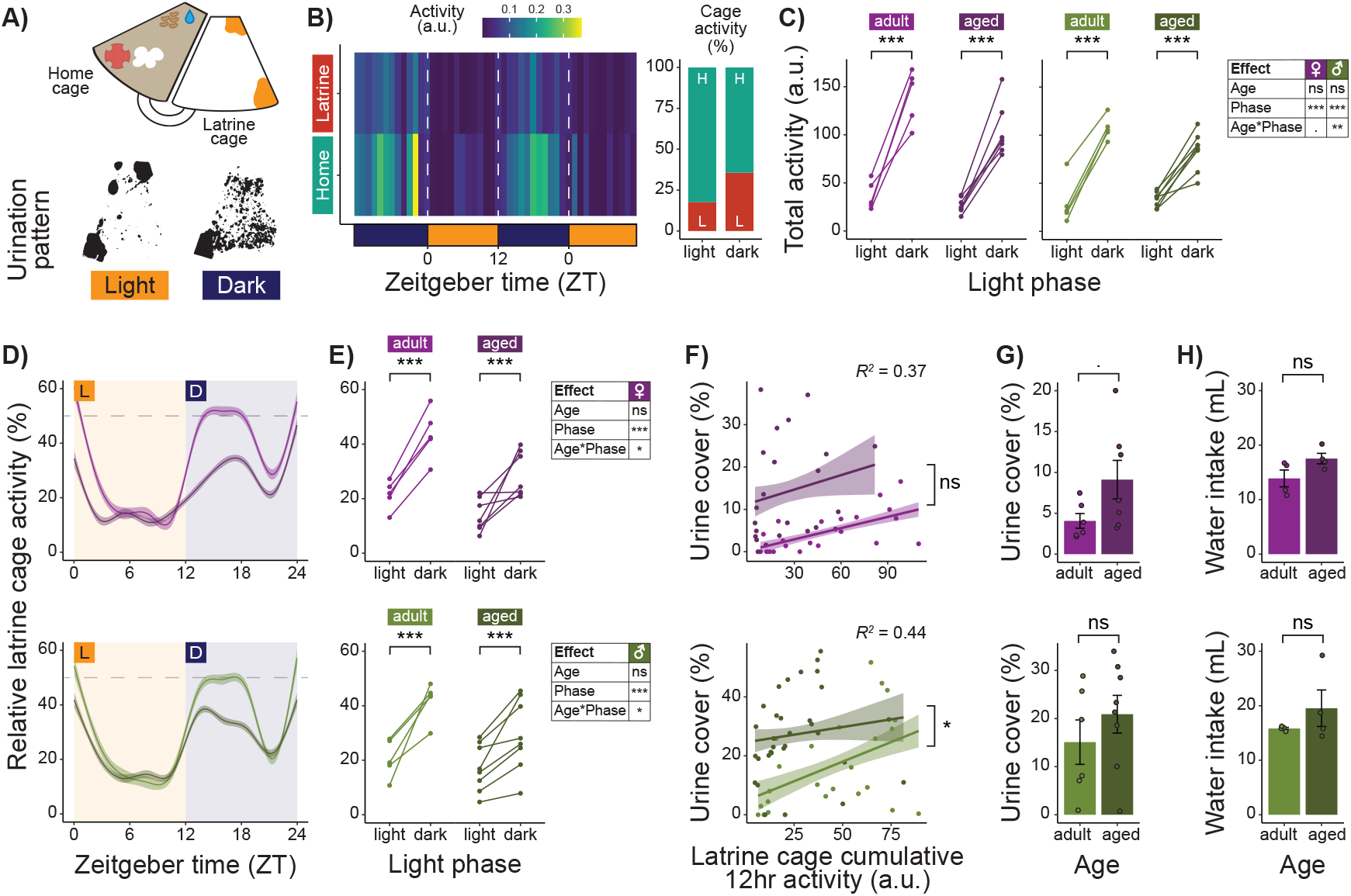
Diurnal regulation of urination behavior in group-housed mice. **A)** Schematic of the latrine cage assay (LCA) complex housing system (top) and representative latrine cage urination patterns (bottom). **B)** Representative 48-hour activity plot (left) and percentage of total activity observed in the home (teal) and latrine (red) cages (right). **C)** Total cumulative 12-hour activity (home + latrine) in adult and aged females (left) and males (right); min-max normalized activity values are expressed in arbitrary units (a.u.); each data point represents the mean of 2 trials per time point per group. **D)** Circadian changes in relative latrine cage activity in adult and aged females (top) and males (bottom) (mean ± SEM). **E)** Average relative latrine cage activity per 12-hour segment; data points represent the mean of 2 trials per group. **F)** Correlation between cumulative 12-hour latrine cage activity and urine cover in females (top) and males (bottom). **G)** Differences in urine cover between adult and aged females (top) and males (bottom); each data point represents the mean of 8 trials per group. **H)** Differences in 24-hour water consumption between adult and aged females (top) and males (bottom); each data point represents the mean of 5 trials per group. Error bars and shaded areas represent SEM. Significance: \*\*\**P* < 0.001; \*\**P* < 0.01; \**P* < 0.05; .*P* < 0.1. Sample size: *n* = 6 adult female, 7 aged female, 6 adult male, and 8 aged male groups.

However, this light/dark activity difference was attenuated in aged mice. While overall activity did not differ between adult and aged mice (**Fig. 2C**; LMMs, female: *t* = 0.630, *P* = 0.535; male: *t*= 0.364, *P* = 0.721) total cumulative activity was 3.2 times higher during the dark phase in adult females compared to 3.0 times higher in aged females (**Fig. 2C**; LMM, age*phase: *t* = 1.766, *P* = 0.086). Similarly, total cumulative activity was 3.5 times higher during the dark phase in adult males but only 1.6 times higher in aged males (**Fig. 2C**; LMM, age*phase: *t* = 3.296, *P* = 0.002). These results indicate reduced diurnal regulation of locomotor activity in aged mice, with a stronger effect in males.

### Diurnal Regulation of Urination Behavior is Dampened in Aged Mice

#### Latrine Cage Assay

To control for variation in overall activity levels among groups, we analyzed the proportion of total activity occurring in the latrine cage on a minute-by-minute basis. Linear regression accounting for temporal autocorrelation revealed a significant age-by-phase interaction in both sexes, indicating differences in circadian expression of relative latrine cage activity between adult and aged mice (**Fig. 2D**; female: *t* = 5.250, *P* < 0.001; male: *t* = 4.984, *P* < 0.001).

Next, we analyzed the average relative latrine cage activity per 12-hour period. LMMs controlling for the number of mice per cage (fixed effect) and cage ID (random effect) confirmed that relative latrine cage activity was higher during the dark phase across all groups (**Fig. 2E**; LMMs, adult female: *t* = 11.573, *P* < 0.001; aged female: *t* = 4.865, *P* < 0.001; adult male: *t* = 8.053, *P* < 0.001; aged male: *t* = 4.763, *P* < 0.001). However, the difference between the light and dark phases was significantly reduced in aged mice (**Fig. 2E**; LMMs, age*phase: females: *t* = 2.061, *P* = 0.047; males: *t* = 2.049, *P* = 0.048). Specifically, relative latrine cage activity was 22.3% higher during the dark phase in adult females but only 14.9% higher in aged females. Similarly, adult males showed 21.3% higher latrine cage activity during the dark phase compared to only 12.0% higher in aged males, representing a reduction of nearly half the circadian amplitude.

Critically, latrine cage cumulative activity was positively correlated with urine cover, confirming that mice were indeed using the latrine cage for urination. LMMs controlling for mouse number and mouse weight (fixed effects), and cage ID (random effect) revealed a significant positive relationship between latrine cage activity and urine cover (**Fig. 2F**; females: marginal R2 = 0.374, *P* = 0.003; males: marginal R2 = 0.441, *P* = 0.011). The slope of this relationship was stronger in adult males than aged males (**Fig. 2F**; LMM, age*activity: *t* = 0.265, *P* = 0.014) but statistically indistinguishable in adult and aged females (**Fig. 2F**; LMM, age*activity: *t* = 1.319, *P* = 0.197). We observed no significant difference in overall urine cover (**Fig. 2G**; LMMs, female: *t* = 1.996, *P* = 0.073; male: *t* = 0.991, *P* = 0.342) or 24-hour water consumption (**Fig. 2H**; LMMs, female: *t*= 1.633, *P* = 0.115; male: *t* = 0.527, *P* = 0.617) between adult and aged mice. Together, these findings suggest that while both age groups use the latrine cage for urination, diurnal regulation of latrine cage activity is significantly reduced in aged mice.

Next, we investigated total urination patterns by temporarily removing the absorptive bedding from the home cage and lining both the home and latrine cages with custom-cut filter paper (**Fig. S2B**). In adult mice, we found that urine cover was significantly higher during the dark phase in both cages (**Fig. S2I**; LMMs, adult female: *t* = 4.004, *P* < 0.001; adult male: *t* = 2.866, *P* = 0.007). However, aged females urinated more during the dark phase only in the latrine cage (**Fig. S2J**; LMMs, home: *t* = 0.705, *P* = 0.497; latrine: *t* = 2.654, *P* = 0.024), and aged males showed increased dark phase urination only in the home cage (**Fig. S2J**; LMMs, home: *t* = 2.321, *P* = 0.049; latrine: *t* = 1.423, *P* = 0.185). Importantly, activity levels in adult mice were positively correlated with urine cover in the latrine cage, but not in the home cage (**Fig. S2K**; LMMs, latrine: *t* = 3.533, *P* = 0.001; home: *t* = 1.345, *P* = 0.186), suggesting that while mice do not urinate exclusively in the latrine cage, they primarily use it for urination, whereas the home cage is used for additional purposes. In contrast, no significant relationship between activity and urine cover was observed in either cage for aged mice (**Fig. S2L**; LMMs, latrine: *t* = 1.256, *P* = 0.224; home: *t* = 0.086, *P* = 0.932). These results indicate that the natural pattern of segregating living and toileting areas—a hallmark of voluntary urination in adult mice—breaks down in aged mice.

#### Void Spot Assay

Next, to quantify individual-level urination behavior, we conducted 90-minute void spot assays (VSAs) during the light (ZT0–ZT3) and dark phases (ZT12–ZT15) (**Fig. 3A**). LMMs controlling for mouse ID and cage ID (nested random effects) revealed that maximum void volume (i.e., volume of the largest void) was significantly larger during the dark phase for all groups (**Fig. 3B**; LMMs, adult female: *t* = 3.972, *P* < 0.001; aged female: *t* = 3.396, *P* = 0.001; adult male: *t* = 4.534, *P* < 0.001; aged male: *t* = 2.903, *P* = 0.005). However, the light/dark difference in maximum void volume was greater in adult males (134 µL on average) compared to aged males (99 µL on average) (**Fig. 3B**). We also compared light/dark differences in total void volume across groups. LMMs showed a significant effect of phase in adult males only (**Fig. 3C**; LMMs, adult female: *t* = 0.117, *P* = 0.907; aged female: *t* = 1.526, *P* = 0.133; adult male: *t* = 2.566, *P* = 0.013; aged male: *t* = 1.831, *P* = 0.071). Finally, aged females exhibited significantly higher phenotypic variance in both maximum and total void volume compared to adults (F tests; maximum void volume: *F* = 0.304, *P* < 0.001; total void volume: *F* = 0.595, *P* = 0.029). These findings demonstrate that aged females show greater variability in urination behavior, both within and among individuals, while aged males exhibit blunted diurnal regulation compared to adult controls.

**Figure 3.**
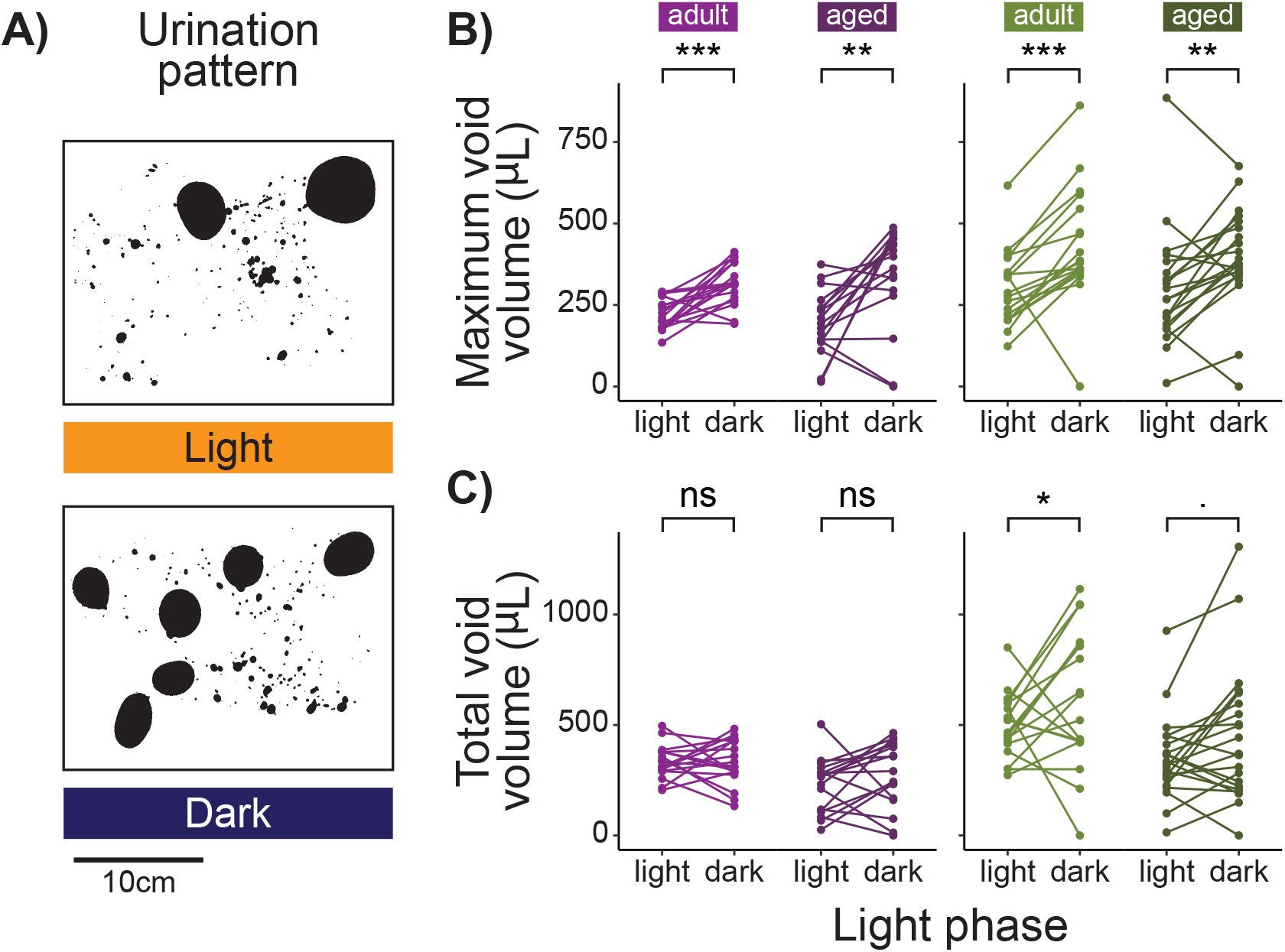
Diurnal regulation of urine volume in individual mice. **A)** Representative light phase (top) and dark phase (bottom) urination patterns in the 90-minute void spot assay (VSA). **B)** Maximum void volume per VSA trial in adult and aged females (left) and males (right); each data point represents the mean of 2 trials per time point per mouse. **C)** Total void volume per VSA trial in adult and aged females (left) and males (right); each data point represents the mean of 2 trials per time point per mouse. Error bars and shaded areas represent SEM. Significance: \*\*\**P* < 0.001;\*\**P* < 0.01; \**P* < 0.05; .*P* < 0.1. Sample size: *n* = 18 adult female, 17 aged female, 18 adult male, and 21 aged male mice.

### Diurnal Gene Expression is Disrupted in Aged Kidney and Bladder

#### Real-time PCR

We hypothesized that urinary system organs (e.g., kidney, bladder) would show aging-related disruption of diurnal gene regulation. We therefore quantified relative transcript abundance of the circadian genes *Bmal*1 and *Per2* in mouse kidneys and bladders collected during the light (ZT0–ZT3) and dark phases (ZT12–ZT15) using real-time PCR (RT-PCR). While adult animals showed diurnal regulation of *Bmal*1 and *Per2* in the kidney (**Fig. 4A**; LMs, adult female *Bmal*1: *t* = 4.159, *P* = 0.002; adult male *Bmal*1: *t* = 3.981, *P* = 0.003; **Fig. 4B**; LMs, adult female *Per2*: *t* = 2.537, *P* = 0.032; adult male *Per2*: *t* = 6.490, *P* < 0.001), aged mice did not (**Fig. 4A**; LMs, aged female *Bmal*1: *t* = 2.026, *P* = 0.078; aged male *Bmal*1: *t* = 0.813, *P* = 0.435; **Fig. 4B**; LMs, aged female *Per2*: *t* = 2.322, *P* = 0.053; aged male *Per2*: *t* = 2.059, *P* = 0.067). These findings suggest that aging disrupts the normal diurnal pattern of circadian gene expression in the kidney.

**Figure 4.**
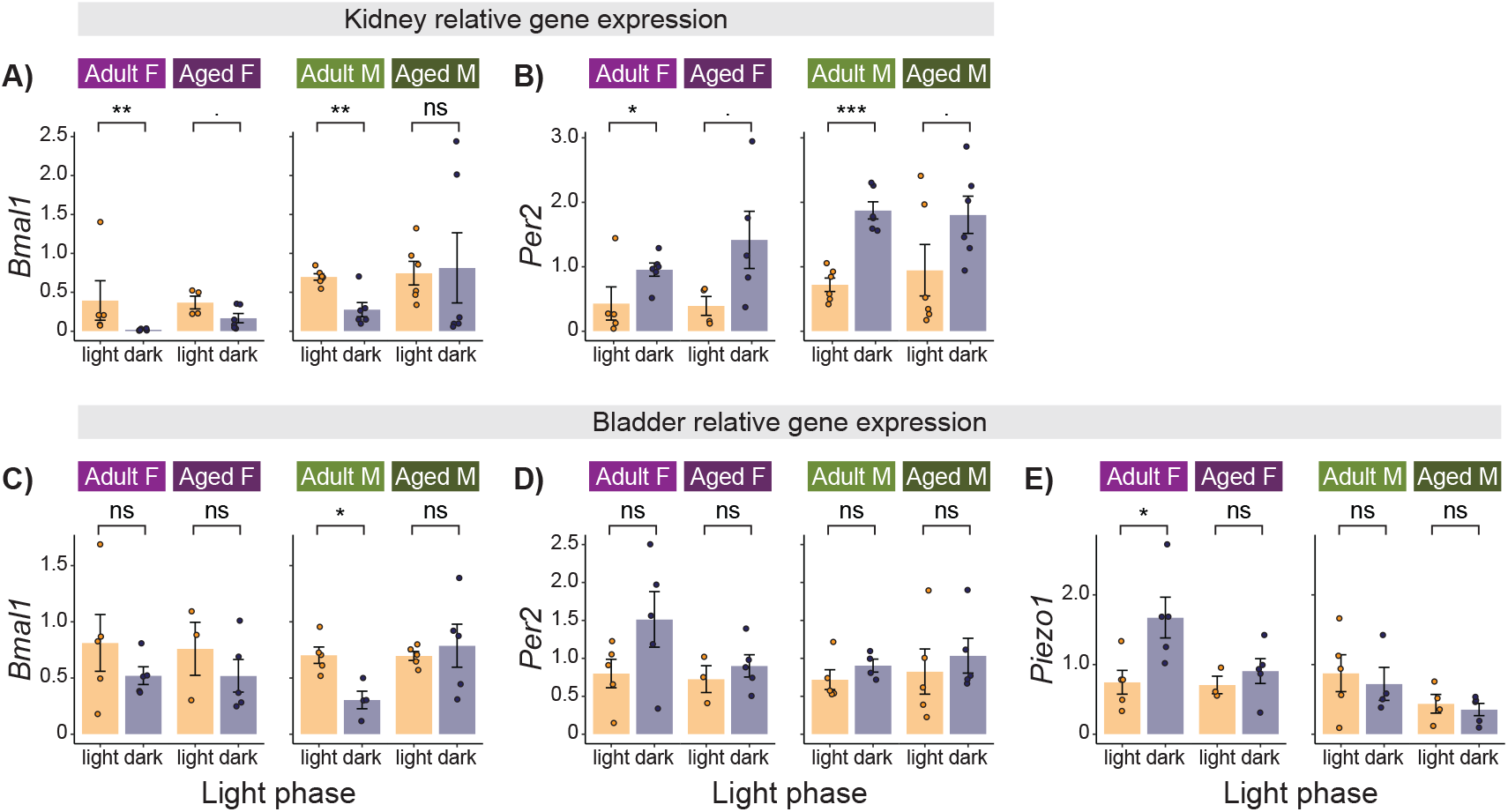
Diurnal expression of core clock genes in the kidney and bladder of adult and aged mice. Relative expression of *Bmal*1 **(A)** and *Per2* **(B)** in the kidney of adult and aged females (left) and males (right). Relative expression of *Bmal*1 **(C)**, *Per2* **(D)**, and *Piezo*1 **(E)** in the bladder of adult and aged females (left) and males (right). Values are normalized relative to the expression of light phase mice of each sex and age group per tissue. Error bars represent SEM. Significance:\*\*\**P* < 0.001; \*\**P* < 0.01; \**P* < 0.05; .*P* < 0.1. Sample size: *n* = 12 adult female, 12 aged female, 12 adult male, and 12 aged male mice.

For *Bmal*1 bladder expression, we detected no effect of age or phase in females (**Fig. 4C**; LM, age: *t* = 0.159, *P* = 0.876; phase: *t* = 1.197, *P* = 0.250). We found significantly higher *Bmal*1 expression during the dark phase compared to the light phase in adult males (**Fig. 4C**; LM, phase: *t* = 3.370, *P* = 0.012), but this effect was absent in aged males (phase: *t* = 0.132, *P* = 0.898). For *Per2* bladder expression, we found no effect of age or light phase in either sex (**Fig. 4D**; LMs, females: age: *t* = 0.959, *P* = 0.353; phase: *t* = 1.675, *P* = 0.115; males: age: *t* = 0.200, *P* = 0.844; phase: *t* = 1.287,*P* = 0.217). These results indicate age-related loss of diurnal regulation of *Bmal*1 in the male bladder.

In mammals, bladder fullness is detected in part by *Piezo* mechanosensitive ion channels (21, 22). Additionally, *Piezo*1 is regulated by clock genes and shows higher expression during the active phase (23, 24). We therefore hypothesized that *Piezo*1 diurnal expression would be disrupted in aged mice. Indeed, we found significantly higher expression during the dark phase compared to the light phase in adult females, whereas this pattern was notably absent in aged females (**Fig. 4E**; LMs, adult female: *t* = 2.936, *P* = 0.019; aged female: *t* = 0.569, *P* = 0.590). We found no significant effect of age or phase in males (**Fig. 4E**; LM, age: *t* = 1.794, *P* = 0.093; phase: *t* = 0.320, *P* = 0.753). Together, these findings suggest that diurnal expression of *Piezo*1 is disrupted in aged female bladders.

#### RNAscope

The urinary bladder is a complex organ composed of diverse cell types with unique transcription profiles (25). To better understand this complexity, we characterized spatial mRNA expression of two circadian genes (*Bmal*1 and *Per*1) and *Piezo*1 using fluorescent in situ hybridization (FISH) in bladder tissue sections. Expression in the detrusor smooth muscle and urothelium was quantified separately (**Fig. 5A, B**). We found no significant expression differences between detrusor and urothelium for *Bmal*1 (LM: *t* = 0.938, *P* = 0.350) or *Per*1 (LM: *t* = 1.066, *P*= 0.288). We therefore combined data from both tissue layers for subsequent analysis.

**Figure 5.**
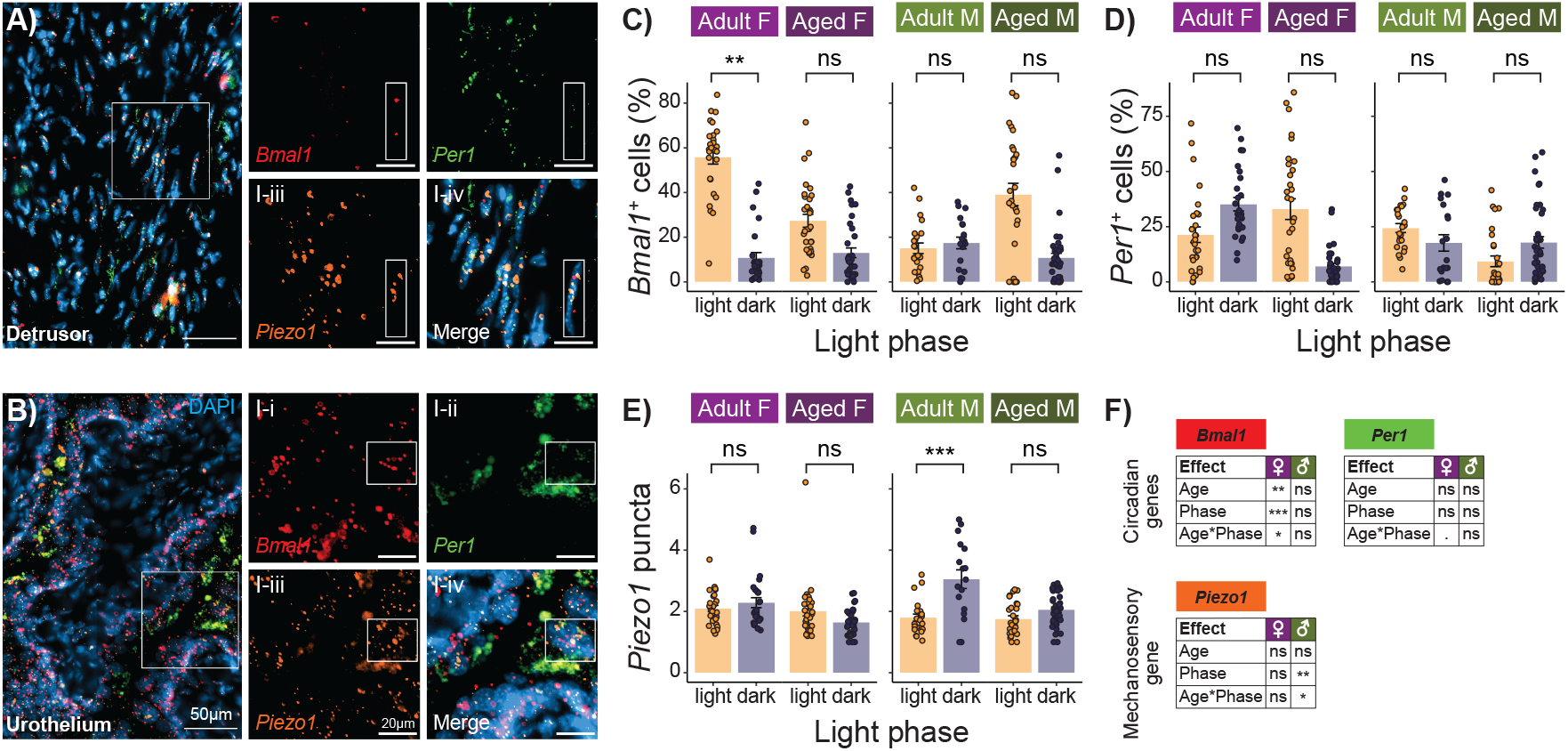
*In situ* hybridization of detrusor and urothelium tissue layers in adult and aged mouse bladders. Representative image of *Bmal*1 (red), *Per*1 (green), and *Piezo*1 (orange) expression in detrusor smooth muscle **(A)** and urothelium **(B)**. Percentage of *Bmal*1 **(C)** and *Per*1**(D)** positive cells (detrusor + urothelium) in adult and aged females (left) and males (right). **E)**Mean number of *Piezo*1 puncta per cell (detrusor + urothelium) in adult and aged females (left)and males (right); each data point represents the mean value per tissue section. **F)** Summary statistic tables for C-E. Error bars represent SEM. Significance: \*\*\**P* < 0.001; \*\**P* < 0.01; \**P* < 0.05; .*P* < 0.1. Sample size: *n* = 6 adult female, 4 aged female, 4 adult male, and 6 aged male mice.

For *Bmal*1, an LMM controlling for mouse ID (random effect) revealed significant effects of age, phase, and their interaction in females (**Fig. 5C**; LMM, age: *t* = 4.801, *P* = 0.005; phase: *t* = 8.212, *P* < 0.001; age*phase: *t* = 3.494, *P* = 0.018). Adult females exhibited a significantly higher proportion of *Bmal*1-positive cells during the light phase compared to the dark phase (light: 56% vs. dark: 11%; LMM: *t* = 7.341, *P* = 0.002), whereas this difference was reduced in aged females (light: 27% vs. dark: 13%; LMM: *t* = 3.412, *P* = 0.190). In contrast, no significant effects of age, phase, or their interaction were observed in males (**Fig. 5C**; LMM, age: *t* = 0.760, *P* = 0.477; phase: *t* = 0.160, *P* = 0.878; age*phase: *t* = 0.703, *P* = 0.510).

For *Per*1, we found a marginally significant age-by-phase interaction in females (**Fig. 5D**; LMM, age: *t* = 0.486, *P* = 0.646; phase: *t* = 1.603, *P* = 0.161; age*phase: *t* = 2.356, *P* = 0.061). Adult females showed a higher proportion of *Per*1-positive cells during the dark phase (light: 21% vs. dark: 35%; LMM: *t* = 1.866, *P* = 0.133), while this pattern was reversed in aged females (light: 33% vs. dark: 7%; LMM: *t* = 1.313, *P* = 0.332). In males, no significant effects of age, phase, or their interaction were observed (**Fig. 5D**; LMM, age: *t* = 1.587, *P* = 0.160; phase: *t* = 0.983, *P* = 0.363; age*phase: *t* = 1.348, *P* = 0.227). Taken together, the expression patterns of *Bmal*1 and *Per*1 in adult females exhibited diurnal regulation consistent with previous findings (26), while these patterns were absent in aged females.

For *Piezo*1, expression was significantly higher in the urothelium than in the detrusor (LM: *t* = 2.052, *P* = 0.041). We therefore included tissue layer as a fixed effect in all LMMs. While the proportion of *Piezo*1-positive cells did not differ between the light and dark phases (LMM: *t* = 0.324, *P* = 0.750), the mean number of *Piezo*1 puncta per cell was significantly higher during the dark phase (LMM: *t* = 2.108, *P* = 0.049). In females, we found no significant effects of age, phase, or their interaction (**Fig. 5E**; LMM, age: *t* = 0.757, *P* = 0.482; phase: *t* = 0.575, *P* = 0.586; age*phase: *t* = 0.619, *P* = 0.561). However, in males, a significant effect of phase and a significant age-by-phase interaction were detected (**Fig. 5E**; LMM, age: *t* = 0.167, *P* = 0.875; phase: *t* = 5.263, *P* = 0.003; age*phase: *t* = 3.101, *P* = 0.031). Adult males exhibited significantly higher *Piezo*1 expression during the dark phase compared to the light phase (LMM: *t* = 4.141, *P* < 0.001), but this light/dark difference was absent in aged males (LMM: *t* = 1.475, *P* = 0.233). These findings suggest that strong diurnal regulation of *Piezo*1 expression in adult males is diminished with aging.

## DISCUSSION

Here, we established a new paradigm in which to investigate the effect of aging on urination at the behavioral and molecular levels. We show that aging disrupts the circadian regulation of urination behavior in mice, accompanied by altered expression of key circadian (*Bmal*1, *Per*1*/2*) and mechanosensory (*Piezo*1) genes in the urinary system. Using a newly developed latrine cage assay, we demonstrate that aged mice exhibit dampened diurnal urination behavior compared to adult controls. Together, these results offer new insights into the pathophysiological mechanisms underlying age-related nocturia and suggest potential therapeutic targets.

Our latrine cage assay represents a significant methodological advance, enabling circadian monitoring of group-housed mice without the confounding effects of social isolation and chronic stress associated with traditional metabolic cages (18). This approach revealed that while overall activity levels remained similar between groups, aged mice of both sexes showed significantly dampened circadian regulation of relative latrine cage activity. This finding parallels clinical observations in older humans, where the normal circadian rhythm of voiding is often disrupted.

Consistent with previous work (27), we observed distinct aging phenotypes between males and females. Aged males exhibited a more pronounced reduction in diurnal regulation of latrine cage activity, with the light/dark difference decreasing by nearly half compared to adult males. This aligns with clinical data showing higher prevalence of age-related nocturia in men than women (3). Additionally, the significant light/dark difference in total void volume observed in adult males in the VSA was absent in aged males. In contrast, aged females showed greater inter-individual variability in VSA urination parameters, including maximum and total void volume. These sex differences may reflect distinct underlying mechanisms of age-related urinary dysfunction and suggest that therapeutic approaches may need to be sex specific.

At the molecular level, our results demonstrate that aging disrupts normal circadian expression patterns of clock genes in the urinary system. The loss of diurnal regulation of *Bmal*1 and *Per2* in aged mouse kidneys parallels the observed behavioral changes, suggesting a mechanistic link between molecular clock disruption and aberrant urination patterns. This finding supports the hypothesis that deterioration of molecular clock function contributes to age-related nocturia (28).

The discovery that normal *Piezo*1 rhythms are disrupted in aged mouse bladders suggests that aging affects both circadian gene expression and mechanosensory pathways in the urinary system. Given the established role of *Piezo*1 in mechanotransduction (22, 29, 30), its circadian dysregulation could contribute to altered bladder sensitivity and inappropriately timed voiding responses in aged individuals. This finding opens new avenues for therapeutic intervention, potentially through targeting mechanosensory pathways in a time-dependent manner.

Our findings have several important clinical implications. First, they support the hypothesis that nocturia should be considered a circadian disorder (28), suggesting that chronotherapeutic approaches might be more effective than current treatments such as bedtime water restriction or antidiuretic administration, which are poorly tolerated in older adults (31). The sex-specific nature of urinary dysfunction in aged mice indicates that therapeutic strategies may need to differ for men and women. Moreover, the disruption of the molecular clock suggests that targeting circadian pathways, perhaps through timed administration of existing medications, could provide new therapeutic opportunities.

Limitations of our study include the inability of our latrine cage assay to track individual urination patterns within groups. Future studies using advanced tracking technologies could help resolve individual contributions to the observed group-level circadian patterns. Additionally, investigation of other circadian-regulated genes and downstream pathways in the urinary system could provide a more complete understanding of the molecular mechanisms underlying age-related nocturia.

Looking forward, our findings suggest several promising research directions. First, determining whether circadian disruption of mechanosensory gene expression in the bladder alters sensory signaling in the brain could help clarify age-related changes in the brain-bladder axis (32). Second, investigating the potential benefits of timing existing medications according to circadian rhythms could lead to immediate clinical applications (33). Finally, developing targeted approaches to restore normal circadian gene expression in the urinary system could provide novel therapeutic strategies for age-related nocturia.

Overall, our study establishes circadian disruption as a key contributor to age-related loss of diurnal urination behavior and provides a new framework for understanding and treating nocturia in the aging population. The sex-specific effects of this disruption and the involvement of both clock genes and mechanosensory pathways suggest multiple potential therapeutic targets for this common condition.

## Funding

This work was supported by an Institutional Development Award (IDeA) from the National Institute of General Medical Sciences of the National Institutes of Health (grant number 2P20GM103432).

## Conflict of Interest

The authors declare no conflicts of interest.

## Data Availability

Data and analysis scripts are available on GitHub (https://github.com/bedford-lab/nocturia).

## Acknowledgements

The authors would like to thank Morgane Vandendoren and Robert Carroll for assistance with mouse husbandry. Research reported in this publication was supported in part by the Institutional Development Awards (IDeA) from the National Institute of General Medical Sciences of the National Institutes of Health under grant number P20GM121310, which supports the University of Wyoming Integrated Microscopy Core.

## Author Contributions

D.S.T, E.E.S, D.R.B, and N.L.B designed the experiments. C.R.W, J.G.L, A.C.N, and E.E.S performed the temperature logger experiments. D.S.T, A.A.A, R.E.F, and N.L.B performed the behavioral experiments. D.S.T, A.C.N, and N.L.B analyzed the behavioral data. S.B.S, S.S.B, and D.R.B performed and analyzed RT-PCR. D.S.T, A.A.A, and R.E.F performed and analyzed RNAscope. D.S.T and N.L.B wrote the manuscript with input from all authors.

## FIGURE LEGENDS

**Supplemental Figure 1.**
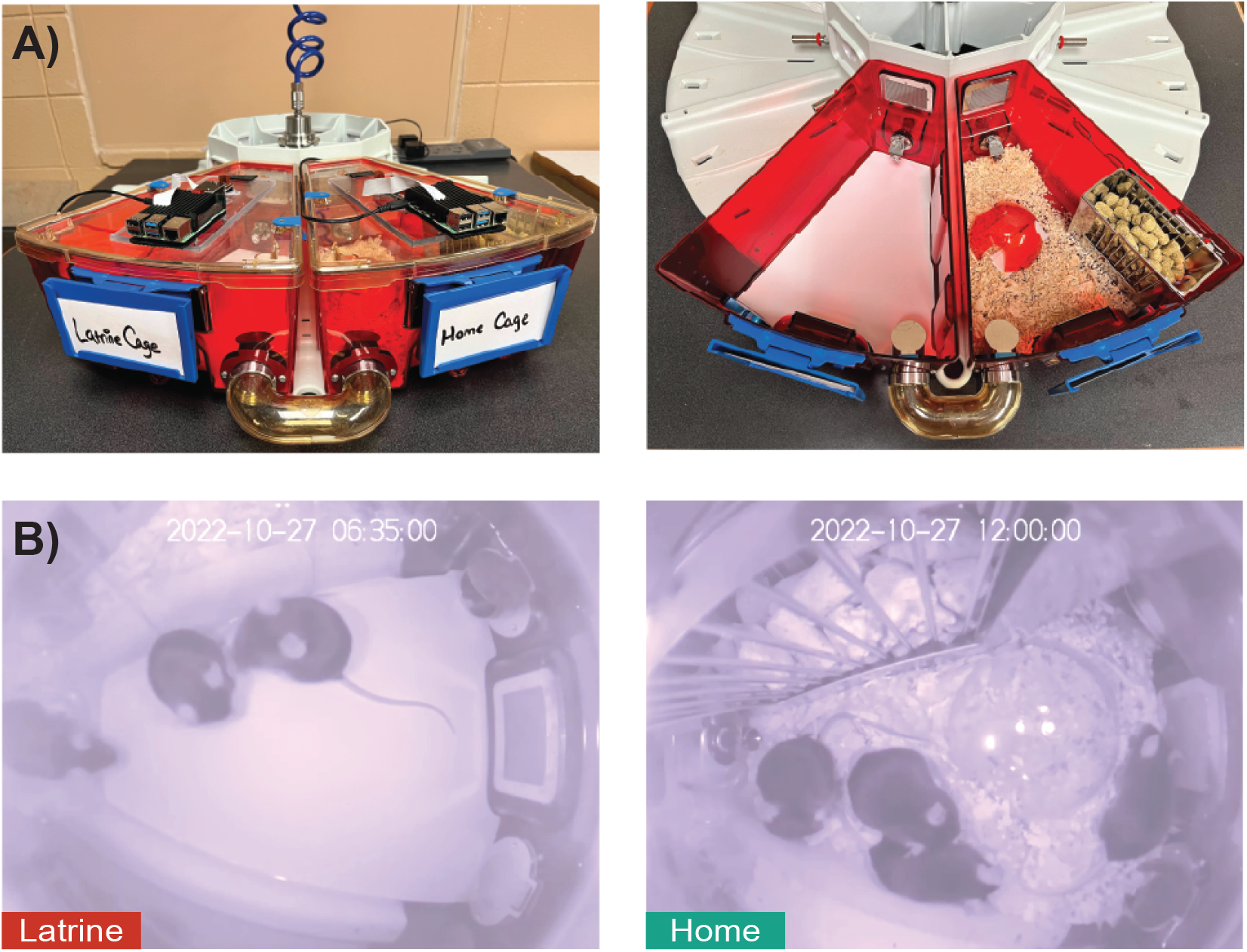
Latrine Cage Assay. The latrine cage (left) is lined with custom-cut filter paper and connected to the home cage (right) via an external tunnel that allows free access between cages **(A)**. Both cage lids are equipped with Raspberry Pi IR cameras and fisheye lenses allowing 48-hour undisturbed recordings of group-housed mice **(B)**.

**Supplemental Figure 2.**
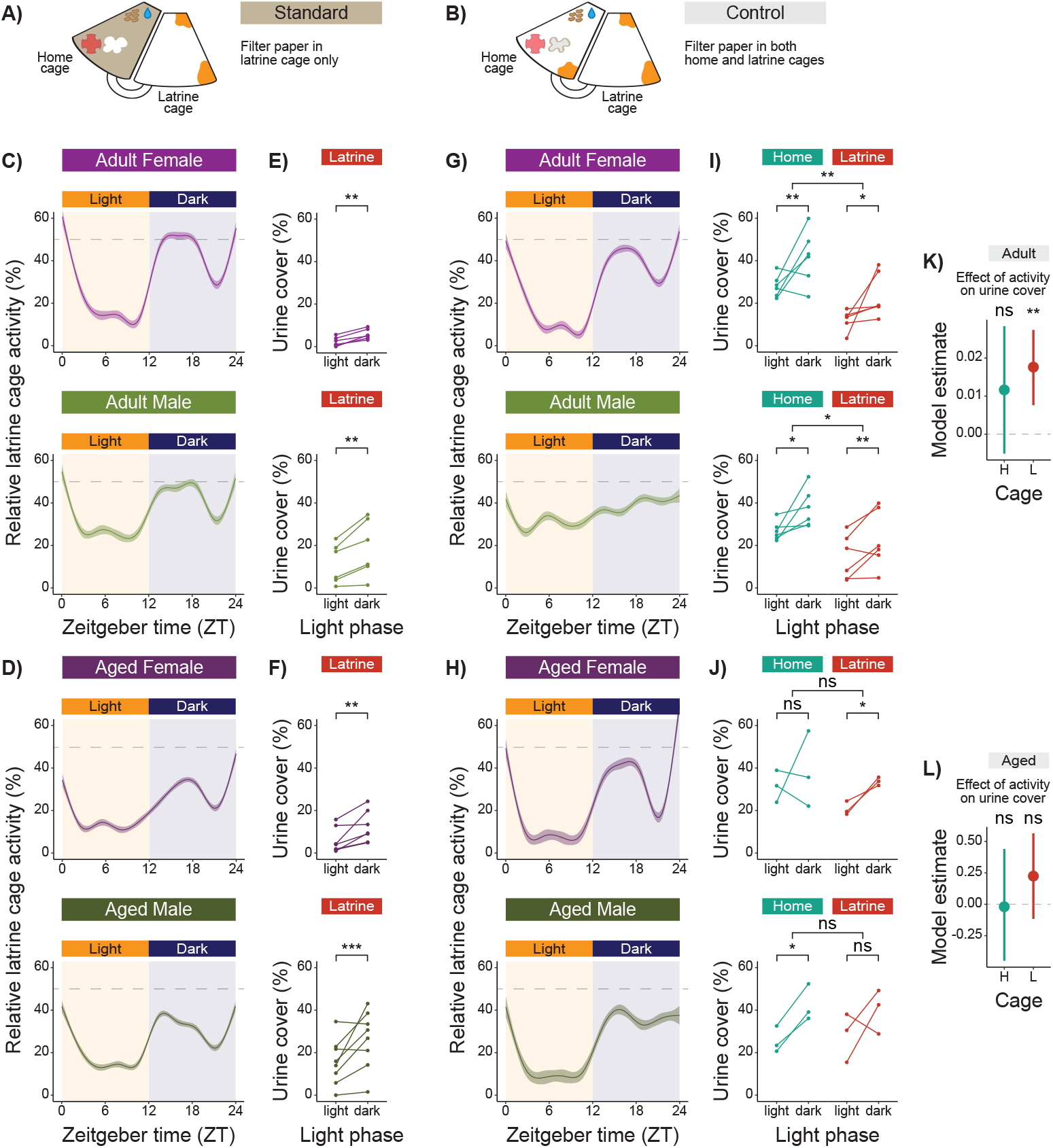
Diurnal regulation of urination behavior in the Latrine Cage Assay. **A)** Schematic of the 48-hour standard latrine cage assay (LCA) with filter paper lining the latrine cage only. **B)** Schematic of the 24-hour control LCA with filter paper lining both home and latrine cages. **C)** Circadian changes in relative latrine cage activity in adult females (top) and males (bottom) in the standard LCA (mean ± SEM). **D)** Circadian changes in relative latrine cage activity in aged females (top) and males (bottom) in the standard LCA (mean ± SEM). **E)** Diurnal differences in latrine cage urine cover in adult females (top) and males (bottom). **F)** Diurnal differences in latrine cage urine cover in aged females (top) and males (bottom). **G)** Circadian changes in relative latrine cage activity in adult females (top) and males (bottom) in the control LCA (mean ± SEM). **H)** Circadian changes in relative latrine cage activity in aged females (top) and males (bottom) in the control LCA (mean ± SEM). **I)** Diurnal differences in home and latrine cage urine cover in adult females (top) and males (bottom). **J)** Diurnal differences in home and latrine cage urine cover in aged females (top) and males (bottom). **K)** Coefficient plot showing regression estimates and 95% confidence intervals for the effect of cumulative 12-hour activity on urine cover in the home (teal) and latrine (red) cages of adult mice. **L)** Coefficient plot showing regression estimates and 95% confidence intervals for the effect of cumulative 12-hour activity on urine cover in the home and latrine cages of aged mice. Significance: \*\*\**P* < 0.001; \*\**P* < 0.01;\**P* < 0.05. Control LCA sample size: *n* = 6 adult female, 3 aged female, 6 adult male, and 3 aged male groups.

**Supplemental Figure 3.**
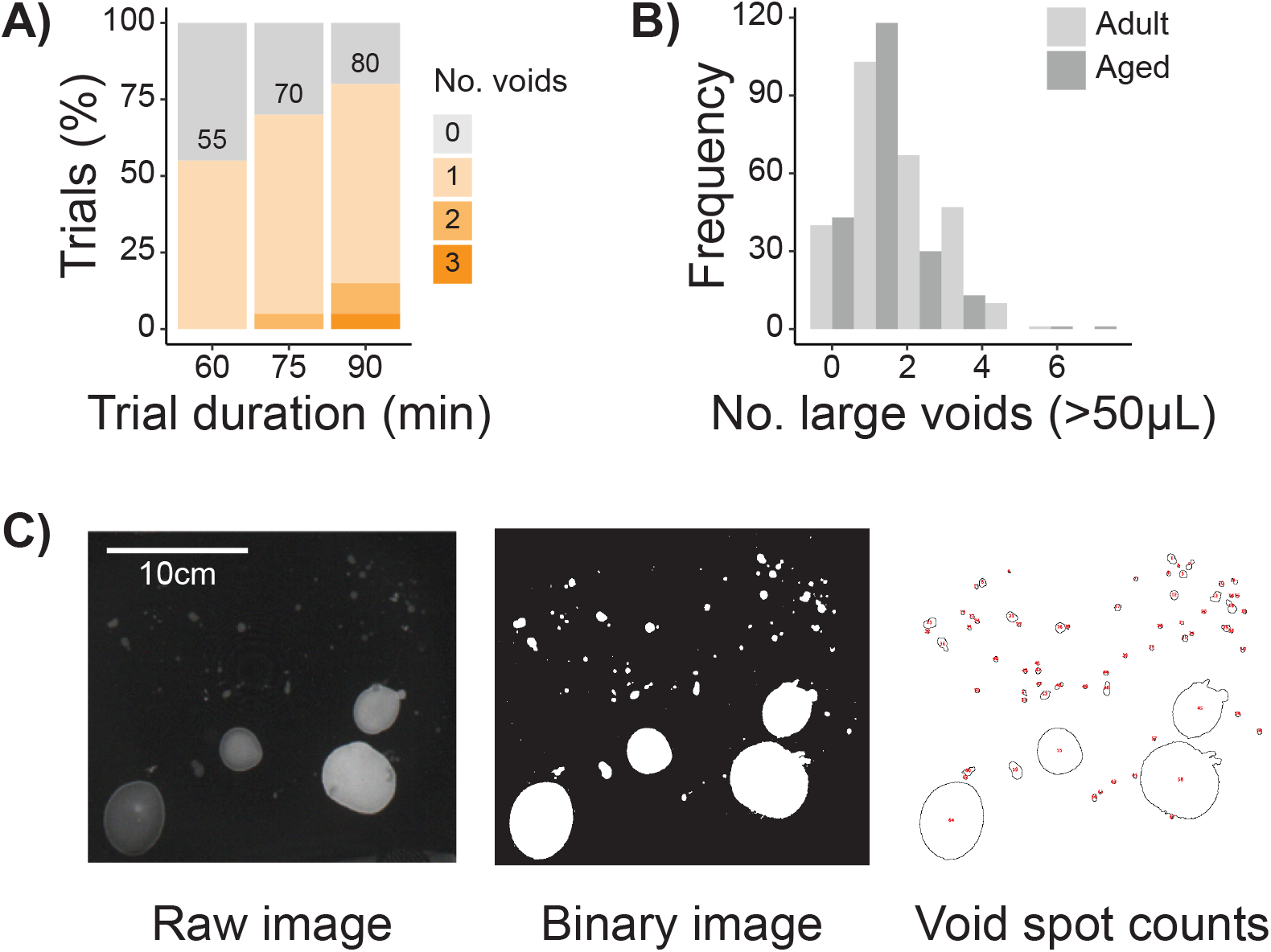
Void Spot Assay. **A)** Number of voids observed during the first 60, 75, and 90 minutes of the void spot assay (VSA) in 20 pilot trials with aged mice. Sample size: *n* = 3 female and 2 aged male mice. **B)** Number of large voids (>50µL) per 90-minute VSA in adult and aged mice (all trials). **C)** Schematic of the ImageJ pipeline for analyzing void spots.

## Notes

### Competing Interest Statement

The authors have declared no competing interest.

### Summary of Updates

Author list and affiliations updated; Acknowledgments updated; Supplemental Figure 2 updated.

