## Supplementary Material for "Diurnal Regulation of Urinary Behavior and Gene Expression in Aged Mice"

### METHODS

#### Animals

Experiments were conducted using adult (4–5 months) and aged (19–21 months) C57BL/6 mice obtained from The Jackson Laboratory (C57BL/6J) and the NIA Aged Rodent Colony. Mice were co-housed in groups of two to five individuals of the same age and sex and were acclimated to the University of Wyoming animal facility for a minimum of two weeks prior to testing. Animals were housed under a 12-hour light:12-hour dark cycle at 22°C in individually ventilated Optimice cages (Animal Care Systems). Cages were furnished with Rocky Mountain softwood screened shavings and quarter-inch corncob bedding (Inotiv). Enrichment included a 2-square-inch cotton nestlet (Ancare) and a red polycarbonate hut (Bio-Serv). Water and Mouse Diet 9F (LabDiet) were provided *ad libitum*. A total of 94 animals were used for this study: 24 adult females, 22 aged females, 24 adult males, and 24 aged males. All experimental protocols were approved by the Institutional Animal Care and Use Committee at the University of Wyoming (protocol numbers 2022-0126 and 2023-0016).

#### Temperature Logger Implants

To monitor the circadian rhythm of core body temperature (Tb), we implanted mice with abdominal temperature loggers. Briefly, mice were anesthetized using either isoflurane or ketamine (100mg/kg) + xylazine (10mg/kg), a small abdominal incision was made, and either a DST nano-T temperature logger (Star-Oddi) or a SubCue data logger (Canadian Analytical Technologies) was implanted intraperitoneally. Animals were allowed to recover for a minimum of one week prior to recording.

#### Behavior

***Latrine Cage Assay (LCA).*** To quantify group-level behavior, co-housed mice were acclimated to the complex housing system for 48 hours prior to recording. An empty cage was lined with custom-cut cellulose chromatography paper (Fisherbrand) and connected to the home cage via an external tunnel (**Fig. S1A**). The filter paper was replaced every 12 hours. To minimize disturbance, the tunnel was removed when all mice were in the home cage, and the filter paper was exchanged in the latrine cage. The tunnel was then quickly reattached to allow free access between cages.

The housing system was then transferred from the standard rack to the home cage recording suite, where cage lids equipped with Raspberry Pi infrared (IR) cameras and fisheye lenses were installed, allowing undisturbed naturalistic observations (1) (**Fig. S1B**). The full guide and code for the Raspberry Pi monitoring software are available on GitHub (https://github.com/cbs-ntcore/pi_monitor). Recordings continued for 48 hours at 30 frames per second (fps), with random assignment of starting times (ZT0 or ZT12). Filter paper exchanges occurred every 12 hours. After testing, the latrine cage was removed, and mice were returned to the standard cage rack.

Using a subset of cages ($n$ = 6 adult female, 3 aged female, 6 adult male, and 3 aged male) we performed a 24-hour control recording with filter paper lining both the latrine and home cages (**Fig. S2B**). Briefly, we removed the absorptive cob bedding from the home cage and installed custom-cut cellulose chromatography paper to quantify urination in the home cage. The home cage still contained food, water, nestlet, and red hut.

***Void Spot Assay (VSA).*** To quantify individual-level urination patterns, we used the void spot assay (2, 3). In a pilot study of 5 aged mice (3 female, 2 male), we conducted 20 trials: 10 during the light phase and 10 during the dark phase. Briefly, animals were placed in an 8” x 10” open field arena lined with cellulose chromatography paper (Fisherbrand) and allowed to freely explore for 90 minutes. We observed at least one void spot within 60 minutes in 55% of trials, within 75 minutes in 70%, and within 90 minutes in 80% (**Fig. S3A**). Based on these findings, we standardized the assay duration to 90 minutes. Light phase VSAs were conducted between ZT0–ZT3, and dark phase VSAs between ZT12–ZT15. Each mouse was tested twice during each phase, with trial order randomized.

***Water consumption.*** We monitored 24-hour water consumption over 3-6 days (mean = 5) for 16 cages (4 adult female, 4 aged female, 4 adult male, and 4 aged male). We determined water intake by comparing water bottle weight at the start and end of each 24-hour period. We also recorded mouse weight, number of mice per cage, and the total mouse weight per cage.

**Gene Expression**

***RT-PCR.*** RNA was extracted from snap-frozen tissues collected during the light (ZT0–ZT3) and dark phase (ZT12–ZT15) using standard TRIzol protocols, followed by reverse transcription using an iScript cDNA synthesis kit (Bio-Rad). RT-PCR was performed with SYBR Green Supermix (Bio-Rad). All samples were run in triplicate. Primer sequences are listed in **Table S1**.

**Supplemental Table 1. RT-PCR Primer Sequences.**

| Gene | Forward Primer | Reverse Primer |
|---|---|---|
| *Bmal1* | 5’-ATCAGCGACTTCATGTCTCC-3’ | 5’-CTCCCTTGCATTCTTGATCC-3’ |
| *Per2* | 5’-GCCAAGTTTGTGGAGTTCCTG-3’ | 5’-CTTGCACCTTGACCAGGTAGG-3’ |
| *Piezo1* | 5’-CGGAACCTGACCTTGACAAC-3’ | 5’-CCAACTGGTGCAGGCTGAC-3’ |
| *Rpl26* | 5’-CGAGTCCAGCGAGAGAAGG-3’ | 5’-GCAGTCTTTAATGAAAGCCGTG-3’ |

***RNAscope.*** Bladders were collected at necropsy and immediately frozen in optimal cutting temperature (OCT) compound, then stored at -80°C. Transverse sections (20µm thick) were obtained using a Leica Cryostat. RNAscope multiplex fluorescent V2 assay (ACD-Bio/Bio-Techne) was performed on tissue sections according to the manufacturer’s protocol. Probes targeting *Piezo1* (cat. no. 500511), *Bmal1* (cat. no. 438741-C2), and *Per1* (cat. no. 438751-C3) were paired with Opal fluorophores. Specifically, *Piezo1* was conjugated with Opal 570 (cat. no. FP1488001K), *Bmal1* with Opal 650 (cat. no. FP1496001KT), and *Per1* with Opal 520 (cat. no. FP1487001KT). Opal fluorophores were reconstituted in 100µL dimethylsulfoxide (DMSO) and diluted in TSA buffer to a final concentration of 1:1000. Following the hybridization and amplification steps, excess liquid was removed from the slides, and 2-4 drops of DAPI were applied for 30 seconds before being gently tapped off. Finally, slides were mounted with ProLong Gold and covered with glass coverslips. Tissue sections were imaged using a ZEISS Axioscan Z1 microscope equipped with a 20X objective. Fluorescence was captured using 50nm (for DAPI), 647nm, and 488nm laser lines. ZEN software (v3.7, Carl Zeiss Microscopy) was used for image optimization, including cropping and adjustment of contrast.

### Data Analysis and Statistics

***Temperature Loggers.*** Core body temperature (Tb) was sampled at 60-minute intervals over a minimum of 8 days. Daily patterns of core Tb were visualized using double-plotted actograms generated using ClockLab software (Actimetrics). To analyze circadian rhythmicity, chi-square periodograms, which decompose the time series into constituent cosine and sine waves, were used to estimate the amplitude and period of circadian rhythms and their deviation from 24 hours. Statistical analyses were performed using linear models in R (R Core Team, 2024).

***Filter Papers.*** Filter papers were imaged using a FLIR USB 3.0 camera (FL3-U3-13Y3M-C). We used blue light excitation (455nm 1150mW Mounted LED, ThorLabs) and a GFP filter set with excitation at 460 ± 30nm (ET460/30x) and emission at 525 ± 50nm (ET525/50m, Chroma). Images were acquired with SpinView software (Teledyne) and converted to a grayscale TIFF before processing in ImageJ (4). We used the "Analyze Particles" function to segment void spots and determine their size and location (**Fig. S3C**). We converted area to volume using a standard curve. Briefly, we pipetted known volumes of 37°C mouse urine (0.5 – 600µL) onto cellulose chromatography paper (Fisherbrand) and determined corresponding void size as described above. Using this standard curve, we multiplied void area ($cm^2$) by 12.772 to obtain void volume (µL). We then used custom R scripts to determine percent cover (latrine cage filter papers) and void volume, number, and total volume (VSA filter papers). For the latrine cage filter papers, we used linear mixed effects models (lmer: lme4 R package) controlling for mouse number and mouse weight (fixed effects) and cage ID (random effect) to test for differences in urine percent cover. For the VSA filter papers, we used LMMs controlling for mouse ID and cage ID (nested random effects) to test for differences in maximum and total void volume, with females and males analyzed independently.

***Home Cage Recording Suite Videos.*** We exported .h264 videos and converted to .mp4 using ffmpeg and analyzed each video at 15 fps in EthoVision XT16 (Noldus). Briefly, we analyzed overall activity per cage based on frame-to-frame pixel changes. A researcher blind to sex and age group applied custom settings to each video to optimize the activity threshold for pixel change quantification. We then took the average pixel change per minute time bin and performed feature scaling using min-max normalization. We aligned the time stamps of the home and latrine cages and calculated relative latrine cage activity on a per minute basis (latrine cage activity / latrine cage + home cage activity). We assessed differences in relative latrine cage activity using time series analysis accounting for temporal autocorrelation (corAR1: nlme R package). We also calculated cumulative activity per cage, per 12-hour period. We tested for differences in cumulative activity using LMMs controlling for the number of mice per cage (fixed effect) and cage ID (random effect). Females and males were analyzed separately.

***RT-PCR.*** QuantStudio 5 Design and Analysis Software (v.1.5.2) was used to determine CT values. The target genes (*Bmal1*, *Per2*, and *Piezo1*) were normalized to the non-circadian housekeeping gene *Rpl26*. Relative gene expression (compared to the light phase control group within each sex/age group) was calculated using the ΔΔCt method for each tissue sample. ΔΔCt values were $\log_{10}$ transformed and compared using linear models in R. Females and males were analyzed separately.

***RNAscope Image Analysis.*** Single-channel images were exported from ZEN software as TIFF files and loaded sequentially into ImageJ (FIJI). These images were stacked and exported as a single, multi-channel TIFF file. The stacked images were then processed using custom ImageJ scripts to isolate detrusor and urothelium regions of interest (ROIs). Cell counting and mRNA puncta quantification were performed using CellProfiler (v4.2.8), downloaded from the official website (https://cellprofiler.org/) (5). Analysis pipelines were developed and optimized separately for detrusor and urothelium ROIs. All analyses were conducted on a Dell OptiPlex 5000 system with a 12th generation Intel® Core™ i7-12700 processor. For each gene target, we calculated the proportion of mRNA-positive cells per ROI and the number of mRNA puncta per cell. We used LMMs controlling for mouse ID (random effect) to test for differences in gene expression, with females and males analyzed independently. All ImageJ and R scripts are available on GitHub (https://github.com/bedford-lab/nocturia).
